# Label-free ghost cytometry for manufacturing of cell therapy products

**DOI:** 10.1101/2023.07.09.545884

**Authors:** Kazuki Teranishi, Keisuke Wagatsuma, Keisuke Toda, Hiroko Nomaru, Yuichi Yanagihashi, Hiroshi Ochiai, Satoru Akai, Emi Mochizuki, Yuuki Onda, Keiji Nakagawa, Keiki Sugimoto, Shinya Takahashi, Hideto Yamaguchi, Sadao Ota

## Abstract

Automation and quality control (QC) are critical in manufacturing safe and effective cell and gene therapy products. However, current QC methods, reliant on molecular staining, pose difficulty in in-line testing and can increase manufacturing costs. Here we demonstrate the potential of using label-free ghost cytometry (LF-GC), a machine learning-driven, multidimensional, high-content, and high-throughput flow cytometry approach, in various stages of the cell therapy manufacturing processes. LF-GC accurately quantified cell count and viability of human peripheral blood mononuclear cells (PBMCs) and identified non-apoptotic live cells and early apoptotic/dead cells in PBMCs, T cells and non-T cells in white blood cells (WBCs), activated T cells and quiescent T cells in PBMCs, and particulate impurities in PBMCs. The data support that LF-GC is a non-destructive label-free cell analytical method that can be used to monitor cell numbers, assess viability, identify specific cell subsets or phenotypic states, and remove impurities during cell therapy manufacturing. Thus, LF-GC holds the potential to enable full automation in the manufacturing of cell therapy products with reduced cost and increased efficiency.

## INTRODUCTION

Manufacturing cell therapy products, including chimeric antigen receptor (CAR) T cells, is a complex and labor-intensive process comprising many steps. To manufacture cellular therapies, cells must undergo several quality control (QC) tests, including assessments of cell viability and the proportion of therapeutic cells in a batch. Importantly, all quality criteria must be satisfied under good manufacturing practice (GMP) conditions. ^1^ As the demand for cellular therapies has increased, efforts to find tools that automate, reduce the complexity, and increase the quality of the manufacturing process have been put at the forefront of cell therapy development programs with the broader goal of improving the consistency of cell therapy products.^2^

In addition to (and complementary to) automated cell therapy systems, there is a need for label-free analytical platforms that can perform deeper cellular characterization for a range of cellular phenotypes that reflect the overall health and therapeutic potential of target cells. To date, several label-free cell characterization techniques based on microscopy or spectroscopy have been reported, including digital holographic microscopy, Raman spectroscopy, and infrared spectroscopy for monitoring of cell viability,^3^ metabolites,^4–6^ and CD4+/CD8+ T cell ratios^7^. Incorporation of machine learning into the label-free analysis can further improve the throughput and accuracy of measurements such as cell purity, cell subtypes,^8,9^ T cell activation state,^9^ and the presence of particulates or impurities.^10^ However, label-free cell characterization approaches using microscopy are limited by low image acquisition rates, intensive and time-consuming data processing, and lack of cell sorting capabilities.

Flow cytometry (FCM) is another approach that can be potentially automated and used in combination with automated cell manufacturing platforms to perform basic QC and phenotypic characterization of cells. FCM-based approaches are advantageous in that they are 1) high-throughput 2) minimize human intervention and 3) reduce the potential variability in cell assessment results. However, QC in cell therapy manufacturing using conventional FCM relies on fluorescent staining of cells with antibodies and/or reagents, which imposes safety concerns and can limit the full automation of the manufacturing process. For example, external fluorescent markers or dyes commonly used in FCM analysis can have a negative impact on cellular functions and may cause toxicity ^11^, limiting the use of the cells in downstream functional characterization assays and more importantly, as therapeutic products. Furthermore, immunofluorescent staining in FCM-based approaches needs to follow Good Manufacturing Practice (GMP) standards, which are often complex and require manual intervention by highly experienced operators. ^12,13^ Consequently, the reliance on fluorescent staining in FCM can lead to increased manufacturing costs and analytical variations, hindering its adoption in automated systems.^14^ Given these limitations, a label-free, high-throughput, cell characterization and QC technique that does not require staining of cells with external markers represents a significant step towards realizing a safe, fully automated, and closed process for manufacturing therapeutic cells.

To this end, we previously reported on label-free ghost cytometry (LF-GC), a technique that leverages high-content cell morphology measures to assess a wide range of functional phenotypes. More specifically, in the LF-GC technique, we produce four different label-free ‘optical signatures’ for individual cells that are reflective of their morphology (**Figure 1**): forward scattering ghost motion imaging (fsGMI), backscattering GMI (bsGMI), diffraction GMI (dGMI), and bright-field GMI (bfGMI) waveforms. These waveforms represent optical signatures for individual cells and can be used, either individually or in combination, to train machine learning classifiers to predict target phenotypes which are defined by using ‘ground truth’ labels during supervised training. Trained classifiers are then used to predict cell subsets without the use of labels in subsequent test analyses.

**Figure 1.**
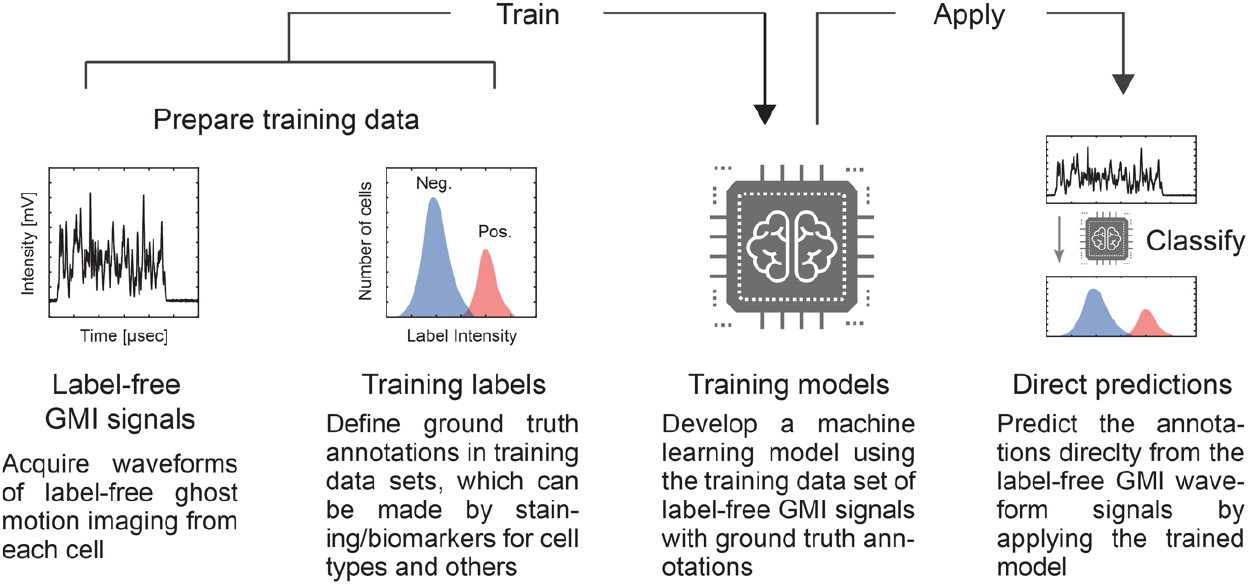
Schematic of a workflow for supervised machine learning in label-free ghost cytometry (LF-GC). A training dataset is prepared from the individual or combination of temporally modulated label-free “imaging” waveforms (GMI: ghost motion “imaging” signals) together with ground truth labels determining the cellular characteristics simultaneously acquired from each cell (left two columns). A machine-learning model is then trained by using the label-free GMI signals annotated with the ground truth labels in the training dataset (the third column from the left). Once the training is complete, specific cell subsets are predicted only by observing the label-free GMI signals (the rightmost column).

LF-GC has been shown to classify many complex phenotypes that are critical for the manufacturing of cellular therapies including; live-dead state, differentiations state, and functional subtyping.^15–17^ In this work, we investigate the use of LF-GC for QC in cell therapy manufacturing. We show that LF-GC is a minimally disruptive, label-free cell analytical method that can be used to characterize important features of cell therapy products including cell number, viability, and phenotypic state. The data suggest that LF-GC is a powerful new approach for monitoring the development of cell therapy products from CAR-T cells to stem-cell-derived therapies.

## RESULTS

### Cell counting by LF-GC

Cell counting (quantifying the total number of cells in a sample excluding debris) is a critical QC step in cell therapy manufacturing. We first assessed the ability of LF-GC to classify cells from debris in human PBMC samples. A classifier model was trained by using the GMI waveforms and acridine orange as a ground truth marker for cells. Acridine orange is a cell-permeant nucleic acid-binding dye that stains all cells, but not debris. After training, the classifier model accurately discriminated cells from debris in PBMCs (ROC-AUC: receiver operating characteristic curves = 0.991, **Figure 2A**). Next, to evaluate the ability of LF-GC to determine cell count, we measured PBMCs at several different concentrations (from 0.25 to 4 × 10^6^ cells/ml). While cell and debris numbers were counted, the sample volume passing through the instrument was measured using a flow meter placed in the sample inlet. The cell concentration of each sample was estimated by the LF-GC-based cell counts and sample volume. As a reference, absolute cell counts were calculated using Flow-Count™ beads spiked into the samples. A linear regression model applied to LF-GC-based cell counts and absolute cell counts showed excellent performance (R^2^ = 0.9996, **Figure 2B**), demonstrating that LF-GC could accurately predict absolute cell counts in human PBMCs at various cell concentrations.

**Figure 2.**
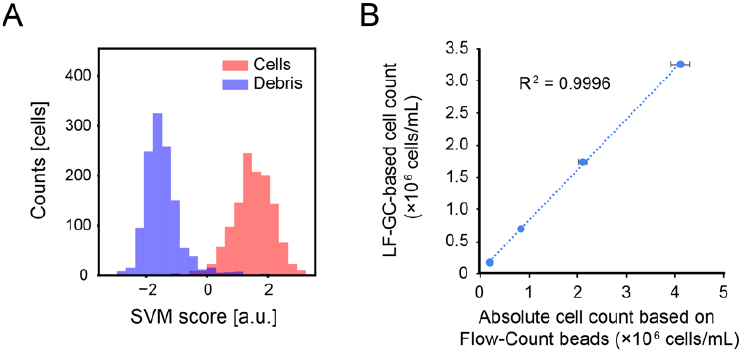
LF-GC-based cell counting in PBMCs. (A) A histogram for the LF-GC-based classification of cells from debris (a.u. = arbitrary units) was obtained by using a support vector machine (SVM) algorithm-based scoring. ROC-AUC for this classification was 0.991. (B) A linear regression analysis between the LF-GC-based cell counts and the absolute cell counts based on Flow-Count beads. The LF-GC-based classification was performed every 10,000 events recorded, and the values represent the average calculated based on 10 independent classifications. Bars indicate SD (standard deviation).

### Viability assessment using LF-GC

Cell viability is directly related to the quality of cell products and accurate quantification of viability is crucial for cell therapy product QC.^18^ To investigate the ability of LF-GC to determine cell viability, we first developed a live/dead cell classifier using human PBMCs. Cell death was induced in PBMCs using 70% ethanol. Live and dead PBMCs were stained with Zombie NIR dye and mixed at different ratios [100:0, 80:20, 50:50, 20:80, 0:100] before LF-GC measurements. We then measured 10,000 events for each sample and pooled the data from 5 samples to train the machine learning model. The model accurately classified live and dead cell populations in a test set of PBMCs (ROC-AUC = 0.9998, **Figure 3A**). Next, we performed live/dead cell classifications on the 5 samples with different live/dead (viability) cell ratios ranging from 0 % to 97.1 %. A linear regression analysis on the LF-GC-based and Zombie NIR dye-based live-cell ratios achieved an excellent fit (R^2^ = 0.9999, **Figure 3B**), showing that LF-GC could measure human PBMC viability with a high degree of accuracy.

**Figure 3.**
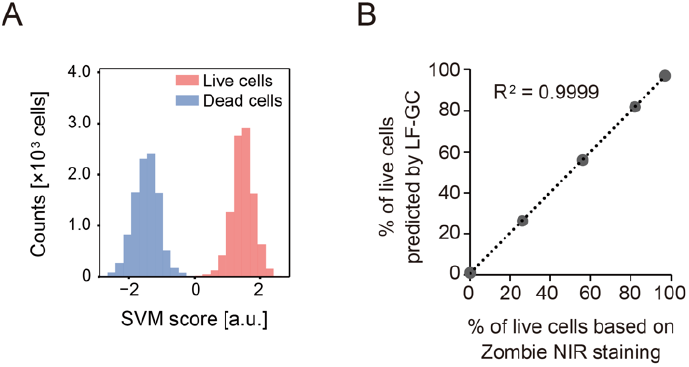
LF-GC-based assessment of live cells in PBMCs. (A) An SVM score histogram for LF-GC-based classification of the live cells from dead cells (a.u. = arbitrary units). ROC-AUC for this classification was 0.991. (B) Simple linear regression analysis between the LF-GC-predicted live cell ratio and the true live cell ratio determined based on Zombie NIR staining. The LF-GC-based classification was performed every 10,000 events recorded, and the values represent the average calculated based on 10 independent classifications. Bars indicate standard deviation.

We next evaluated the ability of LF-GC to detect more complex cell health indicators, which are important to measure for cellular therapy products. For example, apoptotic (or early apoptotic) cells may lack the essential biological functions required for effective cell therapies. In addition, these cells are not expected to proliferate. Thus, early detection of a cell batch with an unusually high proportion of apoptotic cells during the manufacturing process can reduce expenses and increase efficiency. We investigated the ability of LF-GC to discriminate between live and either apoptotic or dead cells in human PBMCs. In the training of a machine learning classifier, we stained cells with a fluorogenic substrate for activated caspase-3/7 (**Figure 4A**) as a ground truth marker for apoptotic cells. We also stained the cells with SytoxAADvanced as a ground truth marker for general viability.

**Figure 4.**
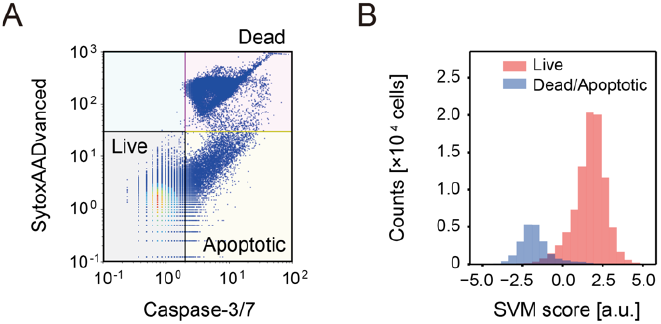
LF-GC-based discrimination of non-apoptotic live cells in PBMCs. (A) Caspase-3/7 activity and SytoxAADvanced scatter plot in which we gated live, apoptotic, and dead cells. (B) An SVM score histogram for the LF-GC-based classification of the live cells and dead/apoptotic cells in PBMCs (a.u. = arbitrary units). The ROC-AUC for this classification was 0.975.

Table 1 shows the performance of the trained classifier in distinguishing between cell populations with different health phenotypes. The model achieved a ROC-AUC score of 0.975 for distinguishing non-apoptotic live cells from apoptotic/dead cells and a ROC-AUC score of 0.911 for differentiating apoptotic cells from non-apoptotic live cells in the test sets. The predicted non-apoptotic live cell percentages (78.1%) closely matched the actual live cell percentages (80.4%) which we define using SytoxAADvanced/activated caspase-3/7. This result supports the ability of LF-GC to accurately distinguish between cells that are alive and in stages of apoptosis using human PBMCs.

**Table 1.** LF-GC-based classification performances live vs. apoptotic/dead cells and live vs. apoptotic cells in PBMCs.

| Live vs. apoptotic/dead cells<br>ROC-AUC | Live vs. apoptotic cells<br>ROC-AUC | Actual live cell rate (%) | Predicted live cell rate (%) |
| --- | --- | --- | --- |
| 0.975 | 0.911 | 80.4 | 78.1 |

### T cell identification and phenotypic analysis using LF-GC

Manufacturing of T cell-based cell therapy products, such as CAR-T cells, requires the accurate measurement of the T cell proportions in a batch, as these can vary throughout their development process. White blood cells (WBCs) from leukapheresis typically serve as the starting material and T cell populations are induced to expand through activation. Since the proportion of T cells increases after activation, the quantification of T cells at different stages of cell therapy production becomes a crucial QC parameter. We, therefore, evaluated the ability of LF-GC to discriminate between T cells and non-T cells in a representative starting material which was WBCs in this study. Fresh blood samples were obtained from a healthy volunteer and subsequently stained with anti-CD3 and anti-CD45 antibodies before being analyzed with LF-GC. The surface marker information of T cells (CD45^+^ CD3^+^) was employed as the ground truth to develop the classifier model (**Figure 5A**). The trained model showed excellent classification performance in detecting T cells from non-T cells in a test set of cell populations with a ROC-AUC of 0.969 (**Figure 5B**).

**Figure 5.**
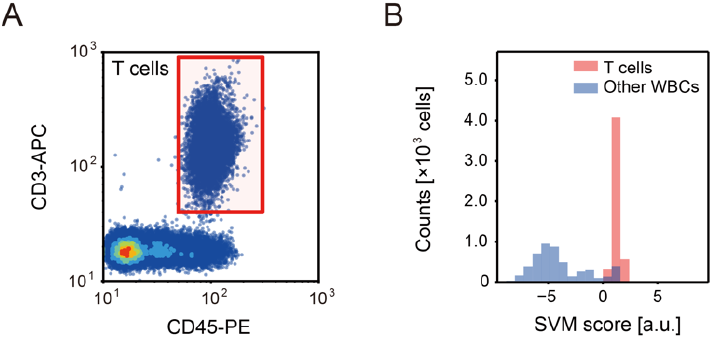
LF-GC-based discrimination of T cells from WBCs. (A) Gating strategy and identification of T cells. (B) SVM score histogram obtained for the LF-GC-based classification of T cells and WBCs (a.u. = arbitrary units). The ROC-AUC for this classification was 0.969.

Next, we assessed the ability of LF-GC to detect more complex T cell phenotypes. During the manufacture of CAR-T cells, T cells are activated before the transduction with CARs. T cell activation is important for both expanding the overall T cell number and increasing the efficiency of CAR transduction, maximizing CAR-T yield. We, therefore, evaluated the ability of LF-GC to differentiate between activated and resting T cells. To prepare populations of activated and resting T cells, after dividing PBMCs from a single donor into two populations, we cultured half with anti-CD3/CD28 antibodies and half without. The two populations were combined on day 5 and subject to LF-GC analysis. Among live cells, the proportions of activated T cells (CD3+/CD25+) and resting T cells (CD3+/CD25-) were 57.5% and 28.0%, respectively. The label-free GMI waveforms obtained from these two populations served as training datasets to develop the classifier models (**Figure 6A**). The trained model showed outstanding classification performance in detecting activated from resting T cells in a test set of cell populations with a ROC-AUC of 0.990 (**Figure 6B**).

**Figure 6.**
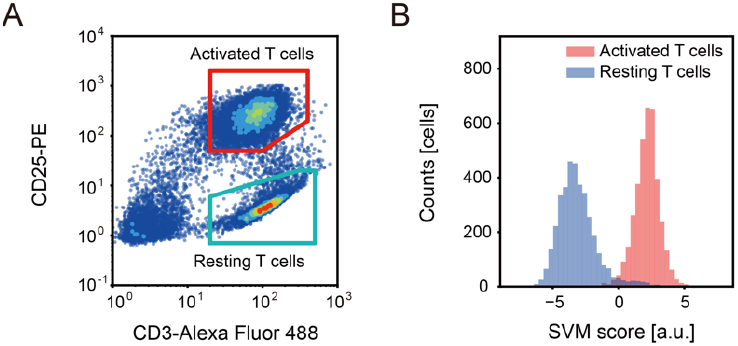
LF-GC-based classification of activated T cells from resting T cells. (A) Gating strategy defining the activated T cells and resting T cells. (B) SVM score histogram obtained for the LF-GC-based classification of activated T cells and resting T cells (a.u. = arbitrary units). The ROC-AUC for this classification was 0.990.

### LF-GC for discrimination of particulates and impurities

A challenge in cell therapy manufacturing is the risk of contamination from various externally introduced impurities, such as microplastic particles, which are often added to cell cultures to ensure proper differentiation and activation of cells. As a medical product, strict guidelines must be adhered to for monitoring potential impurities that could raise safety concerns. To evaluate the ability of LF-GC to identify particulate impurities, we spiked fluorescently labeled plastic microbeads into PBMCs to simulate impurity introduction. We used microbeads with diameters ranging from 0.88 to 31.8 μm, a size range overlapping with PBMCs. Physical separation techniques, such as cell strainers or filters, or more sophisticated techniques such as conventional flow cytometry using simple forward scatter (FSC) signals, are incapable of resolving and separating these populations (**Figure 7A**). In contrast, the trained LF-GC model showed excellent classification performance in differentiating PBMCs from particles in a test set with a ROC-AUC of 0.999 (**Figure 7B**), despite the large overlap of cell/particle size.

**Figure 7.**
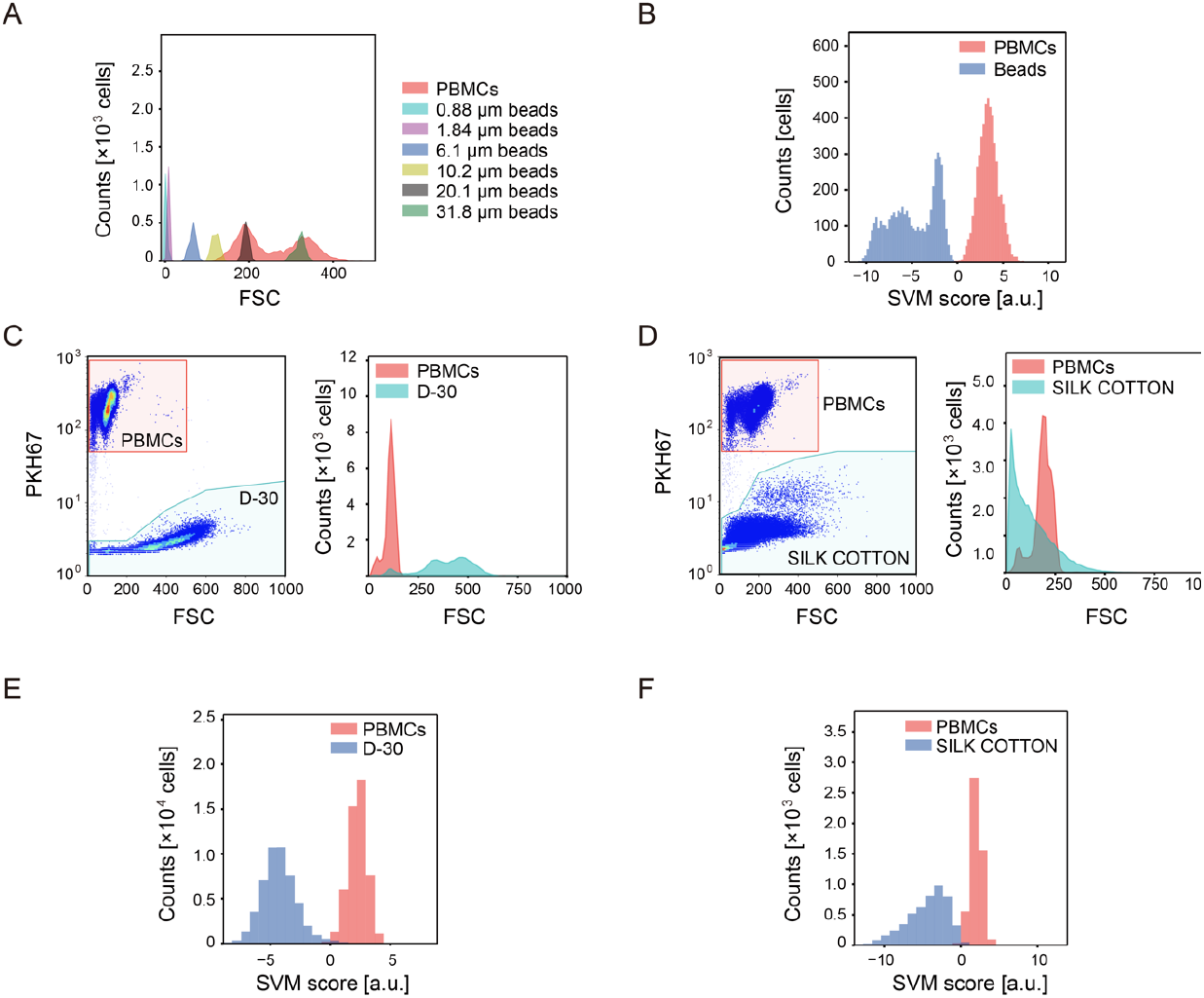
LF-GC-based classification of particle impurities in PBMCs. (A) Size comparison between PBMCs and fluorescent microbeads. (B) SVM score histogram obtained for LF-GC-based classification of PBMCs from beads (a.u. = arbitrary units). The ROC-AUC for this classification exceeded 0.9999. (C) Gating strategy and definition of PBMCs and D-30. (D) Gating strategy and definition of PBMCs and SILK COTTON. Overlays depict comparisons of the FSC intensities from PBMCs and those of D-30 or SILK COTTON. (E) SVM score histogram for the LF-GC-based classification of PBMCs from D-30 (a.u.=arbitrary units). (F) SVM score histogram for the LF-GC-based classification of PBMCs from SILK COTTON (a.u.=arbitrary units). ROC-AUC values for the classification were 0.999 (D-30) and 0.998 (SILK COTTON).

To further validate the use of LF-GC to identify impurities in PBMCs, we spiked cellulose particles, which lack fluorescently distinguishable signatures (D-30 and SILK COTTON), into PBMCs. The shapes of the cellulose particles were different; D-30 is spherical type while SILK COTTON is fiber-like. FSC and BSC profiles of D-30 and SILK COTTON overlap with those of PBMCs, making it difficult to completely distinguish particles from PBMCs based on FSC alone with high accuracy (**Figure 7 C & D**). However, LF-GC separated the cellulose particles well from PBMCs (ROC-AUC = 0.999 for D-30, ROC-AUC = 0.998 for SILK COTTON, **Figure 7 E & F**). These results show that LF-GC can be used to accurately distinguish and will be able to remove a variety of impurities from cell products during the manufacturing process with high accuracy.

## DISCUSSION

Cellular therapies are an emerging class of drugs that have shown particular promise, particularly in the context of oncological conditions.^19^ However, cell therapy manufacturing presents unique and complex challenges that widely span from fundamental biological understanding to quality and regulatory requirements. Large-scale manufacturing of cellular therapies imposes distinct challenges compared to traditional small molecule drugs; manufacturing processes need to be simultaneously developed and refined to increase the yield, batch size, purity, reproducibility, and potency, and all of which have to utilize human cells that will be reintroduced to human patients. These manufacturing challenges have led to the demand for new, minimally invasive tools and technologies to assess cells throughout the drug development process. Although various efforts have been made to lower manufacturing cost through automation and simplification, the complete solution is still elusive; for example, testing for cell types and cell health status still relies on conventional FCM technology with complex and costly procedures.^13,18^

Modern cell therapy manufacturing requires the assessment of multiple characteristics of cell product preparations, from basic cell counts to more complex phenotypic evaluations.^18,20^ Cell counting constitutes a fundamental QC metric needed for the efficient manufacture of cell therapy products.^21^ While FCM can be used for cell counting, its implementation into routine cell therapy manufacturing workflows is hindered by its analytical complexity. To obtain accurate cell counts with FCM, one typically has to add a known concentration of fluorescent beads to samples for absolute cell counting using bead-to-cell ratios and volumetric measurements.^22,23^ We demonstrate that LF-GC accurately counts cells up to 4 × 10^6^ cells/mL and assesses viability up to 97%, which covers the operational range for cell manufacturing. LF-GC thus enables direct measurement of cell lots, without requiring additives such as counting beads or dead cell staining.

Assessing the purity of cell therapy product preparations is another critical aspect. An aim of cell therapy manufacturing is to produce therapeutic cell lots/batches characterized by a high purity, free of exogenous non-biological material. In practice, however, contamination with particulates is an often unavoidable consequence, even in closed systems.^24,25^ Particulate contamination of cell therapies can pose multiple potential safety risks in humans, such as embolism,^24^ tissue inflammation ^26^, or other adverse reactions.^25,27^ Despite the safety concerns, the presence of process-related impurities may not be stringently and comprehensively addressed, as current regulatory standards have been developed and applied primarily for traditional non-cell-based therapies. As the field of cell therapy expands, there is increasing recognition of the need to update regulatory guidance for particulate management.^24^

Besides the challenges of establishing regulations, there are also technical challenges. Current QC approaches for inspecting particulate impurities involve manual microscopy,^21^ a low-throughput, semi-quantitative measurement capable of targeting only a limited range of particulate types. Using LF-GC, we show that particulate impurities could be discriminated with high accuracy, suggesting that LF-GC can be used for the quantitative inspection of particle impurities as well as their removal from culture media. Recently, Grabarek et al.^10^ also reported a label-free method for inspecting particle impurities using flow imaging microscopy combined with deep learning (convolutional neural network). Their observations also support the importance of imaging information and machine learning for the accurate measurement of particle impurities.

Various cellular states, such as healthiness, non-apoptotic conditions, and cell activation can critically influence the quality of cell therapy products. ^28^ Throughout the manufacturing process, apoptosis can be induced by various cellular stresses, such as overgrowth and excessive activation. Moreover, the activation state of T cells significantly affects the gene transduction efficacy of CAR. Traditionally, the detection of apoptotic cells and activated cells has typically employed fluorescent staining methods in FCM-based analysis, leading to increased cost and process complexity. In contrast, this study introduced LF-GC as a potential solution. By accurately discriminating cellular states in a non-destructive, label-free, and high throughput manner, LF-GC provides a more efficient and cost-effective alternative for implementing QC metrics for apoptosis in the cell manufacturing process.

In this study, we focused on testing LF-GC to assess various ‘simple’ phenotypes, such as cell type and viability state, as commonly employed QC metrics. However, we foresee the importance of evaluating the application of LF-GC for the assessment of more complex phenotypes related to the therapeutic potential of cellular therapies. For instance, we have recently shown that LF-GC can accurately classify the exhaustion states of T cells,^17^ of which were reported as a key indicator of CAR-T cell functionality.^28,29^ LF-GC performed 5-part cell type classification from peripheral WBC and classification of polarized M1/M2 macrophages.^16,17^ Lastly, future investigations exploring the use of LF-GC to assess non-T cell therapies such as CAR-macrophages or CAR-NK cells may offer additional insights into any shared or unique mechanisms driving the therapeutic potential of cell-based immunotherapies.

We presented, in this study, the utilization of models trained with supervised machine learning for evaluating T cell therapy products. In this approach, the accuracy of the model can partially depend on whether ground truth labels are assigned to each LF GMI waveform. Since we often mix the training samples and flow them to generate data sets for training models, there were cases where we could not perfectly assign the correct labels to the waveforms. For instance, in Fig 7, the ground truth labels for PBMCs and microparticle populations were defined by gating using a scatter plot of fluorescent intensity of PKH67 and FSC intensity. It is possible that debris originating from PBMC samples and comparable in size to some of the microparticles while being PKH67 negative, may have been erroneously included in the ‘microparticle’ training population. Due to the potential impurity in the training data sets, the classification accuracy of PBMCs and particles obtained in this study could have been compromised. A possible approach to overcome this issue is the employment of an unsupervised machine learning approach, where training populations are defined solely by the morphological differences present in a mixed population.^17^ Future studies that assess a variety of machine learning approaches, including supervised and unsupervised methods as well as their combinations, to define ‘ground truth’ populations will clarify the degree of achievable classification accuracy.

In summary, we show that LF-GC is a promising non-destructive and label-free analytical platform for rapidly and accurately monitoring key QC characteristics of cells during the manufacture of cell therapy products. The versatility of LF-GC permits its application as a part of an automated analytical platform. We foresee that the future development of LF-GC-based in-line cell analyzers and label-free cell sorters specifically designed for cell manufacturing will lead to enhancements in process optimization, reduction in manufacturing costs, and creation of higher quality cellular therapies.

## MATERIALS AND METHODS

### In silico-labeled Ghost Cytometry (LF-GC)

Morphological information of the cells was acquired based on a label-free compressive ghost imaging technique described previously.^15,16^ While full image reconstruction was demonstrated either with iterative acquisition using differently structured illuminations or structured masks with a single-pixel detector,^30,31^ LF-GC adopts a reduced acquisition without image production using machine learning models for performing high-speed cell classification.^16^ All photomultiplier tubes (PMTs) used in this work were purchased from Hamamatsu Photonics Inc. PMTs of 10 MHz with built-in amplifiers (H10723-210 MOD, MOD2, H10723-Y2, A2, MOD2) were used for detecting the dGMI, ssGMI, bsGMI, and fsGMI signals, while PMTs of 200 kHz (H10723-20 Y1, H10723-20 MOD, H10723-210 Y1), 1MHz (H10723-210 MOD3, H10723-20 MOD3), or 10MHz (H10723-210 MOD2) were used to detect fluorescence signals and BSC signals. FSC signals were obtained using a PMT of 200 kHz (H10723-20 MOD, H10723-20-01). Multi-pixel photon counter (MPPC, S13360-6075CS) from Hamamatsu Photonics Inc was used to detect the bfGMI. The direct current of dGMI, fsGMI, bfGMI, ssGMI, and FSC signals was cut with an electronic high-pass filter. The PMT signals were recorded with electronic filters using a digitizer (M2i.4932-Exp, Spectrum, Germany) or an FPGA development board (TR4, Terasic) with a homemade analog/digital converter. The digitizer and/or FPGA continually collected a fixed length of signal segments from each color channel at the same time, with a fixed trigger condition applied to the FSC signals. The dataset containing label-free signals from each cell and the simultaneously acquired fluorescent signals were randomized and split into training and testing datasets. In the training dataset, label-free signals of unique populations of cells and beads were grouped based on fluorescence labels as the ground truth. An SVM algorithm was employed to train a binary classification model that predicts the identity of cells, and an SVM score (decision function) was computed based on label-free signals alone. The performance of the classifier was evaluated using a testing dataset (**Figure 1**); scores of the area under the ROC-AUC, as well as SVM histograms, were considered for this purpose. Training and testing of the model algorithms were performed using either equal amounts of samples for each class or with the class ratios of the samples. SVM score (decision function) histograms described in the figures were derived from the trials with the best AUC.

### Cell counting by LF-GC

Human PBMCs was purchased from Precision for Medicine. Four samples with cell concentrations of 0.25, 1, 2, and 4 × 10^6^ cells/mL were prepared in a FACS buffer containing 2% fetal bovine serum (FBS) and 1 mM EDTA in PBS. The PBMCs were stained with Cellstain Acridine Orange (AO) (Dojindo) at a concentration of 1 μg/mL for 20 min to identify cells from debris. Flow-Count Fluorospheres (Beckman Coulter) with a known particle concentration were added at a 1:1 volume ratio to each sample as a reference for calculating the absolute count of cells in a sample measured by LF-GC.

For the flow cytometric analysis, we used backscatter and AO fluorescence signals to identify Flow-Count Fluorospheres (BSC-high/AO-positive), PBMCs (BSC-med/AO-high), and the rest of singlets as debris. We randomly selected and pooled 10,000 events (5,000 PBMCs and 5,000 debris) from the data acquired for four samples with different concentrations of PBMCs, and trained an SVM-based classifier using this dataset. To assess the classifier’s performance, an additional 1,250 events from each class (2,500 events in total) were selected randomly, ensuring there was no overlap with the training dataset.

For the LF-GC-based cell counts, we predicted the counts of cells for each sample with different cell concentrations (0.25, 1, 2, and 4 × 10^6^ cells/mL) using the above classifier based on SVM (PBMCs vs. debris). We tested the classifier 10 times for each sample with different cell concentrations without overlap and the mean and standard deviation were calculated. The absolute counts were also calculated 10 times for each sample with different cell concentrations, and the mean value was used for the assessment of the performance of LF-GC-based cell count.

### Discrimination of live and dead cells

Frozen human PBMCs were purchased from STEMCELL Technologies, and dead PBMCs were prepared by treating them with 70% ethanol for 30 min, followed by washing with the FACS buffer. Live and dead PBMCs were stained with Zombie NIR dye at a 1:1000 dilution for 20 minutes on ice and, after washing once with the FACS buffer, they were mixed at different ratios (100:0, 80:20, 50:50, 20:80, and 0:100) and analyzed by LF-GC. Among a total of 50,000 events (10,000 events from each condition), we identified debris, live, or dead cells based on FSC, BSC, and Zombie-NIR signals. We trained the SVM methods to classify live cells and dead cells using 5,000 events from each class (10,000 cells in total), selected randomly from the acquired data. We assessed the trained model to discriminate the viability of cells using additional 10,000 events from each class, which were further selected randomly without overlap with the training dataset. The test was repeated 10 times using ten sets of 10,000 cells for each sample, and the average and standard deviation were calculated for each sample.

### Discrimination of non-apoptotic live cells from and apoptotic/dead cells

Frozen human PBMCs were purchased from STEMCELL Technologies. After thawing by immediately placing them into a 37°C water bath and transferring them into a pre-warmed RPMI1640 with 10% FBS and 1% Penicillin-Streptomycin, cells were stained with either the FITC Annexin V Apoptosis Detection Kit (BioLegend) or CellEvent™ Caspase-3/7 Green Flow Cytometry Assay Kit (Thermo Fisher Scientific) to label apoptotic cells (Caspase-3/7 activity-positive and SytoxAADvanced-negative) and dead cells (Caspase-3/7 activity-positive and SytoxAADvanced-positive) according to manufacturer instructions and analyzed using LF-GC.

After splitting the acquired data into two subsets for training or testing SVM-based models, we randomly selected either a set of 3,000 live cells and 3,000 apoptotic/dead cells or another set of 3,000 live cells and 3,000 apoptotic cells from one subset for training data. For the assessment of the classifier’s performance, 89,860 live cells and 21,943 apoptotic/dead cells were used for classifying live and apoptotic/dead cells. 3,000 live cells and 3,000 apoptotic cells were used for classifying live and apoptotic cells.

### Blood sampling and discrimination of T cells from WBCs

All experiments using human blood samples were performed according to ethical standards; written consents were obtained from the blood donors. The study protocol was approved in advance by the Institutional Review Board of the Chiyoda Paramedical Care Clinic (Tokyo, Japan) (Approval No. 15000088) and was registered in the UMIN Clinical Trials Registry (UMIN000045011). Peripheral blood samples were obtained from healthy donors who provided written informed consent. This study was conducted in accordance with the principles of the Declaration of Helsinki. Inclusion and exclusion criteria are as follows:

Inclusion criteria

- Healthy male/female subjects ranging in age from 20 to 59, at informed consent
- Subjects who can give informed consent to take part in this research, after being provided with an explanation of the protocol detail

Exclusion criteria

- Subjects being under some kind of continuous medical treatment, except applying a nasal spray and eye drops
- Subjects who took a steroid within the last three months from the time point of clinic visiting
- Subjects who apply a steroid all the year around, except seasonal use and deteriorating conditions.
- Subjects who were vaccinated within the past two weeks to the clinic visiting.
- Pregnant, possibly pregnant, and lactating women.
- Subjects with previous and/or current medical history of serious diseases in heart, liver, kidney and/or digestive organs.
- Subjects using excessive alcohol.
- Subjects with serious drug and/or food allergies.
- Subjects who are under the other clinical tests with some kind of medicine/food, or participated in those within four weeks after this research, or are planning to join those after giving informed consent to participate in this research.
- Subjects who donated their blood components and/or whole blood (200 mL) within a month to this research.
- Males who donated blood (400mL or more) within the last three months of this research.
- Females who donated blood (400 mL or more) within the last four months of this research.
- Males who will be collected in total of their blood (1200 mL) within the last twelve months, after adding the blood amounts planning to be sampled in this research.
- Females who will be collected in total of their blood (800 mL) within the last twelve months, after adding the blood amounts planning to be sampled in this research.
- Others who have been determined as ineligible for participation, judging from the principal/sub investigator’s opinions.

Fresh blood samples were collected from healthy volunteers in tubes containing EDTA 2 K tubes (BD Biosciences), WBCs were prepared by removing red blood cells using Lysing Buffer (BD Biosciences). WBCs were then stained with PE anti-human CD45 antibody (Thermo Fisher Scientific, 12-0459-42) and APC anti-human CD3 antibody (BioLegend, 300412) for 20 min, washed with FACS buffer, and stained with DAPI (Dojindo) at 1 μg/mL for 5 min. The cells were then analyzed using LF-GC. Doublets, dead cells, and debris were removed from the dataset using the FSC, BSC, and DAPI gates, and the remaining events were used for further analysis. The cells that were double positive for CD45 and CD3 were defined as T cells, while other live cells were defined as non-T cells among WBCs.

We trained an SVM-based classifier using randomly selected 10,000 cells in total (5,000 T cells and 5,000 non-T cells for each class) from the acquired data. To assess the classifier’s performance, we randomly selected an additional 10,000 events in total (5,000 T cells and 5,000 non-T cells for each class) without overlap.

### Discrimination of particle impurities

Human PBMCs were purchased from STEMCELL Technologies and Precision for Medicine. Cellulose particles (without fluorescent labels) and polystyrene particles (with fluorescent labels) were used as impurities, and one of them was spiked into the PBMCs kept in each measurement tube. Two types of cellulose particles [CELLULOBEADS D-30 (D-30) and SILK COTTON PW (SILK COTTON)] were purchased from DAITO KASEI KOGYO CO., LTD. In the experiment using PBMCs spiked with cellulose particles, PBMCs were pre-stained with the PKH67 Green Fluorescent Cell Linker Kit (Merck) as a ground truth label. Zombie NIR Fixable Viability Kit (BioLegend) staining was performed to exclude dead cells and debris from the analysis. All fluorescent polystyrene microbeads were purchased from Spherotech Inc. Six different sizes of fluorescent beads (0.88 μm; FP0852-2 1.84 μm; FH2052-2, 6.1 μm; FP6052-2, 10.2 μm; FP10052-2, 20.1 μm; FP20052-5, and 31.8 μm; FP30052-5) were spiked into PBMCs and analyzed using LF-GC. In the analysis, the acquired data were split into training/testing datasets (for cellulose particles with PBMCs: train size = 24,000, test size = 10,000; for fluorescent plastic microbeads: train size = 7,362, test size = 9,324). The SVM model-based classifier was trained and the classifier’s performance was assessed using equal numbers of datasets of cells and beads.

### Discrimination of activated T cells from resting T cells

Frozen human PBMCs were purchased from Precision for Medicine and were thawed by immediately placing them into a 37°C water bath and transferring them into a pre-warmed RPMI1640 with 10% FBS and 1% Penicillin-Streptomycin. Then, PBMCs were cultured at 37°C under 5% CO_2_ for 5 days in ImmunoCult™-XF T Cell Expansion Medium (STEM CELL Technologies) supplemented with 30 ng/mL human recombinant IL-2 (Peprotech) in the presence or absence of plate-bound anti-CD3 antibody (clone UCHT1, Beckman Coulter) and soluble anti-CD28 antibody (clone CD28.2, BD Biosciences). After 5 days of culture, CD3/CD28-treated PBMCs were mixed with untreated PBMCs. The mixture was then stained with Alexa Fluor 488 anti-human CD3 antibody (BioLegend, 300320), PE anti-human CD25 antibody (BioLegend, 302606), and Zombie NIR™ (BioLegend) for 20 min, washed with FACS buffer, and analyzed using LF-GC. The cells were gated using FSC-height and FSC-width scatter plots to remove doublets, then they were gated using the FSC and BSC scatter plots to remove debris, and gated using the BSC-Zombie NIR scatter plot to remove the dead cells. The gated cells were further gated and labeled as activated T cells (CD3+/CD25+) or resting T cells (CD3+/CD25-). The data was split into two subsets for training/testing datasets. Within these data, 7,400 and 7,400 cells selected randomly (without overlap) with equal numbers of activated T cells and resting T cells were used for training the SVM-based classifier and assessing the classifier’s performance, respectively.

## ACKNOWLEDGEMENTS

We would like to thank all members of ThinkCyte Inc. for experimental support and discussion, in particular, Kaori Matsuyama and Ryusuke Miyazaki for the technical support in sample preparation. We also thank Willem Westra and Hiroaki Adachi for reviewing the manuscript and for the helpful discussion. This study was supported by the Product Commercialization Alliance (PCA) program funded by the New Energy and Industrial Technology Development Organization (NEDO).

## AUTHOR CONTRIBUTIONS

K.T., K.W., H.O., K.S., K.N., S.T., H.Y., and S.O. contributed to the study conception and design. K.T., K.W., K.T., H.N., H.O., S.A., E.M., Y.O., and K.N. acquired the data. H.O., K.T., K.W., Y.Y., and K.N. analyzed and interpreted data. H.O., K.T., K.W., Y.Y., K.S., K.N., and S.O. drafted and revised the manuscript. All the authors approved the final manuscript.

